# Magnetic Temporal Interference For Noninvasive, High-resolution, and Localized Deep Brain Stimulation: Concept Validation

**DOI:** 10.1101/2020.07.20.212845

**Authors:** Mohsen Zaeimbashi, Adam Khalifa, Cunzheng Dong, Yuyi Wei, Sydney Cash, Nian Sun

**Affiliations:** Department of Electrical and Computer Engineering, Northeastern University, Boston, MA; Department of Neurology, Massachusetts General Hospital, Harvard Medical School, Boston, MA

## Abstract

Non-invasive deep brain stimulation has been a major challenge in the field of neuroscience and brain stimulation in the past three decades. Current brain stimulation technologies suffer from such hurdles as the inability to do deep brain stimulation, poor spatial resolution, and invasiveness. Transcranial magnetic stimulation (TMS) technique, for instance, cannot target brain regions deeper than ∼2cm and has a poor spatial resolution, impacting a large area of the peripheral region and leading to various side effects. Implantable electrodes, even though effective for deep brain stimulation, are invasive and carry various drawbacks related to the surgery and site infection. In this paper, we propose a new concept that relies on temporal interference of two high- frequency magnetic fields generated by two electromagnetic coils. The neural system does not respond to each of these high-frequency magnetic fields alone because of the intrinsic low-pass filtering properties of the neural membrane. The peripheral areas of the brain are impacted only by the high-frequency magnetic fields that cannot stimulate the nerves, while the deep brain area where the two fields interfere experiences a magnetic field that contains a low-frequency envelope and therefore the nerves can be stimulated. This technique can noninvasively focus a magnetic or electric beam at any depth inside the brain with a high resolution, without impacting the peripheral regions.

## 1 Introduction

The clinical use of Transcranial Magnetic Stimulation (TMS) is among the most prominent achievements in the field of neuroscience in the past two decades. The use of TMS to intervene with malfunctioning brain circuits has revolutionized the way neural disorders are treated and understood. TMS, as an FDA-approved technique, has provided remarkable relief to patients with Parkinson’s disease, essential termer, dystonia, and other debilitating disorders. The ability to modulate the activity of neural networks noninvasively and painlessly allows researchers to treat disease with much more ease than previous neuromodulation techniques such as deep brain stimulation (DBS). Unfortunately, unlike DBS, TMS is unable to stimulate deep regions of the brain. This is because of the rapid attenuation of the magnetic field generated by the TMS coil, leading to a maximum effective stimulation depth of around 2-3cm beneath the scalp [1, 2]. The magnetic field is weak beyond this range and cannot target the central part of the brain. This is unfortunate as it has been demonstrated, in recent years, that the regions involved in most neuropsychiatric disorders are placed also in non-superficial brain areas and, therefore, the stimulation of these deep structures could be a more efficient treatment of patients with Parkinson’s disease, essential tremor, and dystonia [3, 4]. Despite recent efforts that led to the developed of new coil designs (new shape and size) to improve the focality and stimulation depth, TMS still shows poor spatial resolution and neuromodulation is mainly limited to superficial cortical brain sites [5, 6, 7, 8]. This is caused by the large size of conventional TMS coils, which in addition to target sites, also impacts neighboring brain regions and leads to side effects such as headache, twitching of facial muscles, or lightheadedness [9]. Any development of TMS coils that allow for stimulation beyond the resolution of “figure-of-eight” coils while maintaining the field intensity required to activate neurons will give researchers more opportunities to stimulate specific neural circuits that are important in neurological diseases.

In this work, we propose a new approach for noninvasive, high-resolution, and focalized deep brain stimulation that relies on temporal interference of two high frequency magnetic fields generated by two electromagnetic coils. The neural system does not respond to each of these high frequency magnetic fields alone because of the intrinsic low-pass filtering properties of neural membrane [10]. In other words, brain cannot follow or react to the high frequency magnetic fields (e.g. >1kHz) created by the coils. However, if these two magnetic fields differ by a small amount Δf in frequency, the resulted temporally interfered signal contains a low frequency envelope and the neural system might follow and respond to this envelope.

It is noteworthy that the electrical temporal interference technique is already investigated in [11], where the authors use two pairs of electrodes to directly inject high frequency currents to the brain. However, this technique still suffers from certain limitations due to the need to inject direct current flow through the skin or brain; the authors in this work have proposed two mechanisms to deploy this technique on human brain: 1) placing the electrodes on the skin outside the brain, in which case—because of the low electrical conductivity of the skull—most of the current applied to the electrodes will flow through the skin without actually penetrating through the brain and reaching deep brain area; 2) placing the electrodes under the skull and on the surface of the brain, in which case this technique would still be invasive and require surgery and electrode implantation. In the event that one needs to move the focal point inside the brain and stimulate a different area, relocating the electrodes and further surgery might be required as well.

## 2 Summary and concept of magnetic temporal interference technique

In this work, we propose using two temporally-interfered, high-frequency magnetic fields generated by two electromagnetic coils in order to achieve noninvasive, high-resolution, and focalized deep brain stimulation. The neural system does not respond to each of these high frequency magnetic fields alone because of the intrinsic low-pass filtering properties of neural membrane [10]. I.e. brain cannot follow or react to the high frequency magnetic fields (e.g. >1kHz) created by the coils. However, if these two magnetic fields differ by a small amount Δf in frequency, the resulted temporally interfered signal contains a low frequency envelope and the neural system may follow and respond to this envelope. In this technique, the peripheral area of the brain is impacted only by the high frequency fields *B*_1_(*ω*_1_) and *B*_2_(*ω*_1_ + Δ*f*) that cannot stimulate the nerves, while the deep brain area— where the two fields interfere and create a magnetic field that contains a low frequency envelope Δf—can be stimulated. The focal point where the two fields interfere can be adjusted at any depth inside the brain simply by changing the value of the currents injected to the coils or changing the location of the coils. The schematic of a simulated brain phantom with two MTI coils is shown in Fig. 1. The results show that the field arising from two coils can interfere at the central part of the brain and target deep brain area without impacting the peripheral regions.

**Fig. 1.**
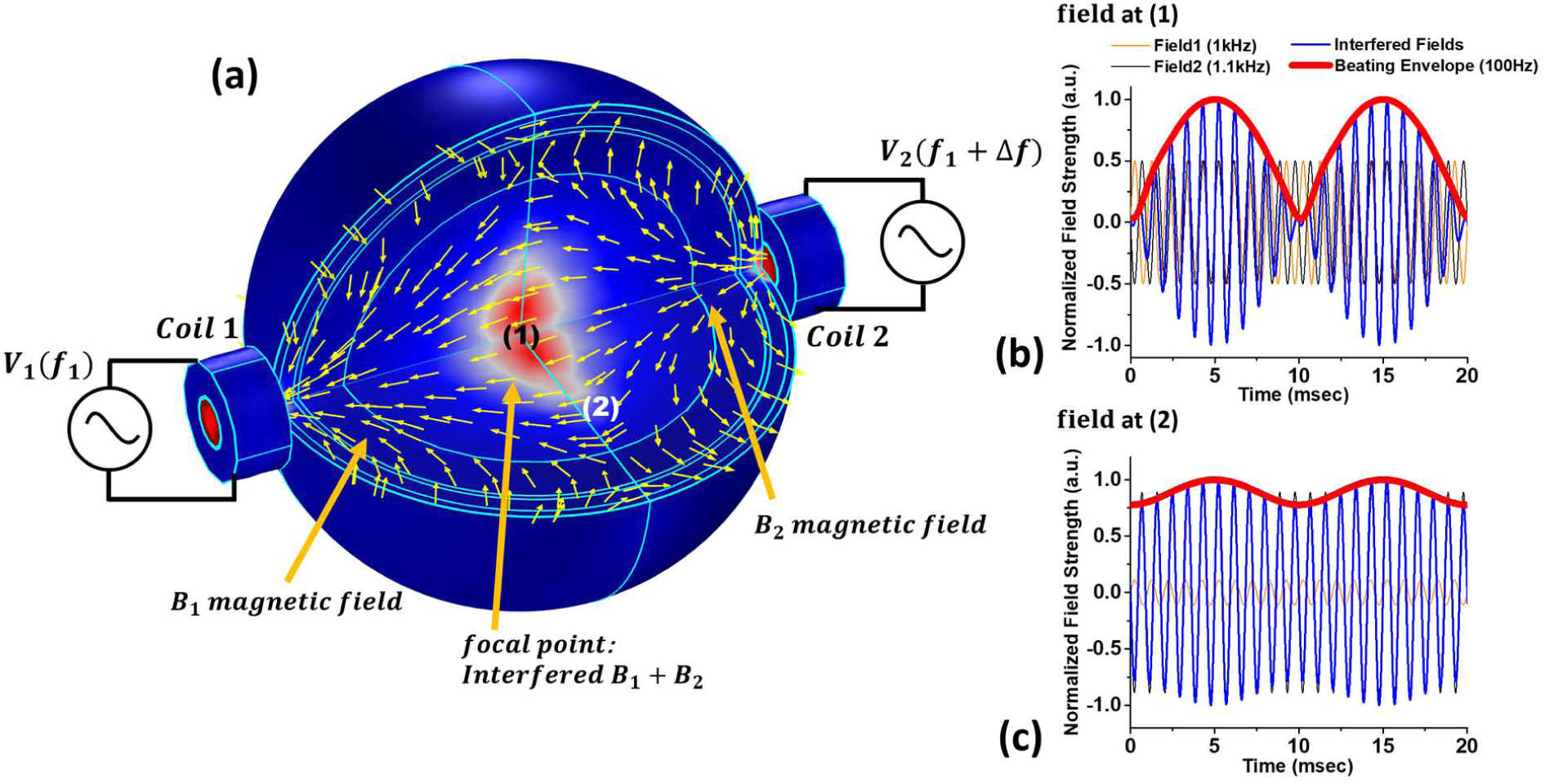
Simulation results showing the concept of MTI on a human brain model with radius of 10cm and different layers including skin, skull, CSF, grey matter, and white matter. (a) Focalized magnetic field beam in deep brain area without impacting the peripheral regions. Coil 1 and coil 2 are excited with a high frequency Sine wave signal at 1kHz and 1.1kHz. The low frequency envelope is the difference between the two frequencies and equal to 100Hz; b) The magnetic field waveform at region (1) in central region, where the two fields interfere and create the low frequency envelope with high amplitude; c) the magnetic field waveform at region (2) outside the central area, showing the low frequency envelope with a small amplitude. The peripheral regions are only impacted by high frequency field 1 and 2 and therefore neurons cannot be stimulated, whereas the central region is impacted by a high-amplitude low frequency envelope and can be stimulated.

## 3 Design, simulation, and concept validation of magnetic temporal interference technique

We have designed, simulated, and optimized the temporal interference technique and coils using COMSOL Multiphysics software. The size of the coils, number of turns, their magnetic core, and locations with respect to the brain are optimized in a way that fulfil multiple objectives: a) the size of the coils and their locations with respect to the brain should be optimized such that the produced magnetic fields by two coils can interfere at any area and any depth inside the brain with a high spatial resolution. Unlike TMS technique, which can only stimulate the peripheral regions and lacks a high spatial resolution, MTI technique can focus the magnetic or electric fields at deep brain area; b) the generated electric field gradient, which is the most important parameter for magnetic brain stimulation, should be higher than the threshold value required for brain stimulation, which is about 11k V/m^2^ [12, 13]; c) the designed coils should be able to efficiently operate at frequencies up to 50kHz. This is because the induced electric field of the coils increases linearly by increasing the operational frequency, meaning that we can generate very large electric fields and go above required threshold value by increasing the frequency and without increasing the current, heavily reducing the generated heat and power consumption. There might be a maximum limit in terms of career frequency beyond which this technique may not be effective enough. This means that we cannot increase the career frequencies indefinitely, even if we keep the low frequency envelope at 100Hz. This matter should be investigated in the future animal studies.

In order to deliver the maximum amount of magnetic and electric fields in the near field region, the coils were designed such that the quality factor 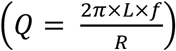 is maximized while satisfying two main constraints: 1) the maximum delivered current that the wire can handle, and 2) the maximum voltage the power amplifier can deliver. The diameter of the coil was first chosen to achieve the desired focal stimulation in a rat brain model. To significantly boost the generated magnetic and electric fields, we added a magnetic core made of MnZn, from Fair-Rite Products Corporation, with a with relative permeability of 17. The MnZn ferrite core can operate at frequencies up to 100kHz (much larger than our operating frequency). Using the optimized parameters shown in Table 1, the coil achieves a Q of 322.

**Table 1.**
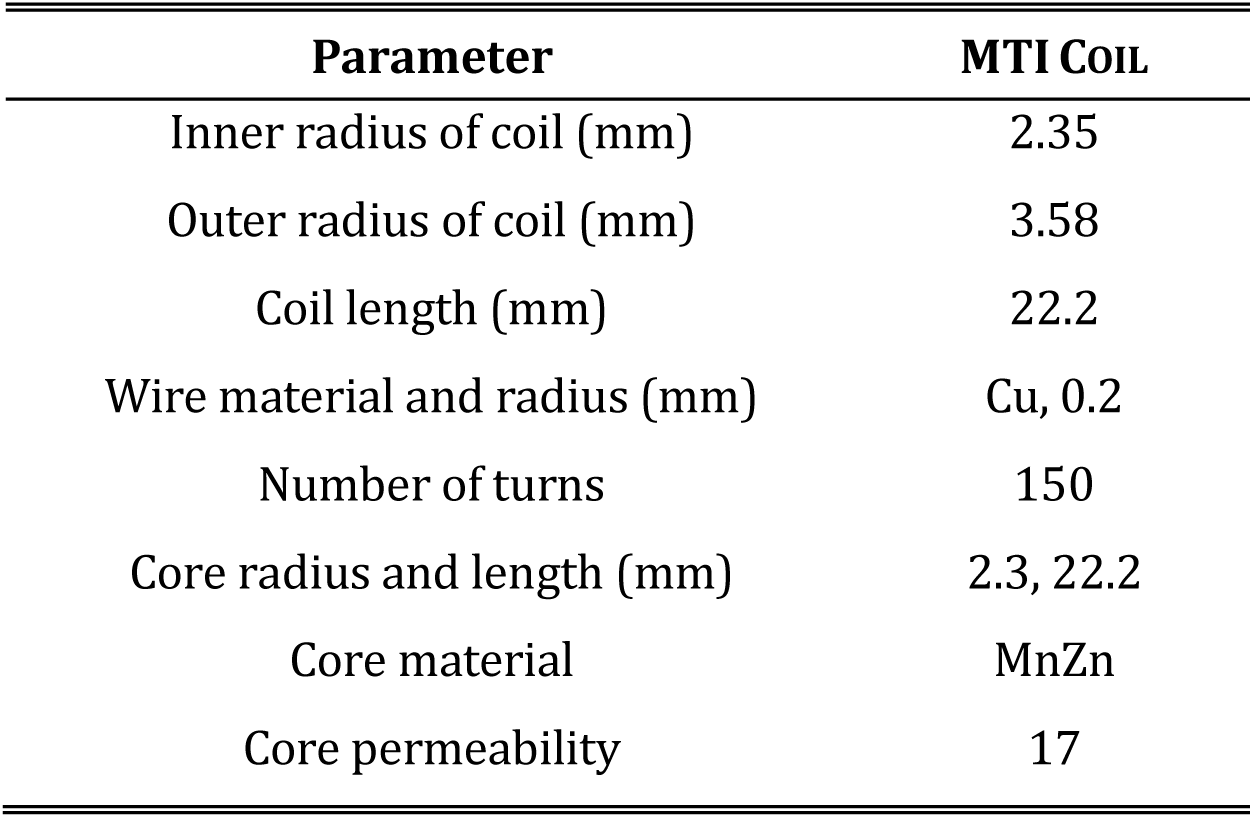
Coil design parameters.

### Simulation data on concept validation of magnetic temporal interference technique

We have simulated the magnetic temporal interference technique in COMSOL and the simulation data shows that this technique can be effective for noninvasive, high-resolution, and localized deep brain stimulation, something that TMS technique cannot do. We have used a 3D rat brain model with the size of 10×16×21 mm^3^ in the simulations. The two designed MTI coils produce a magnetic field, the magnetic field penetrates through the brain and induces electric field and electric field gradient, which can stimulate the brain neural system. The rat model and coil configuration are shown in Fig. 2. As mentioned before, we have used MnZn ferrite with relative permeability of 17 as magnetic core which can significantly boost the generated magnetic field. The pulsed-Sine wave current applied to the coils is 40Amp which runs only for 10 msec instead of continuous powering, reducing the heating effect significantly. The 10msec timing is equivalent to one full cycle of a low frequency envelope of 100Hz. The currents applied to the coils should be at 180° phase difference in order to have full destructive interference at t=0 and full constructive inference at t=5msec. Fig. 3a shows the magnetic field distribution inside the cross-sectioned rat brain when only coil 1 with 50kHz frequency is ON. As it is shown the left part of the brain is impacted by this high frequency signal, but no stimulation is expected since the frequency is too high and neurons cannot respond or follow the generated field [10]. Fig. 3c shows a scenario where only coil 2 with 50.1kHz frequency is ON. The same situation exists here, and neurons cannot respond to this high frequency field and therefore no stimulation happens. Fig. 3e shows the temporally interfered magnetic field distribution inside the cross-sectioned rat brain when both coil 1 and 2 are ON. This field distribution shows the amplitude of the low frequency envelope (100Hz in this case,) which is maximum in central part of the brain and minimum in peripheral region. Neural system in deep brain region can demodulate and respond to this low frequency envelope, which is formed as a result of temporal interference of the two fields from coil 1 and 2.

**Fig. 2.**
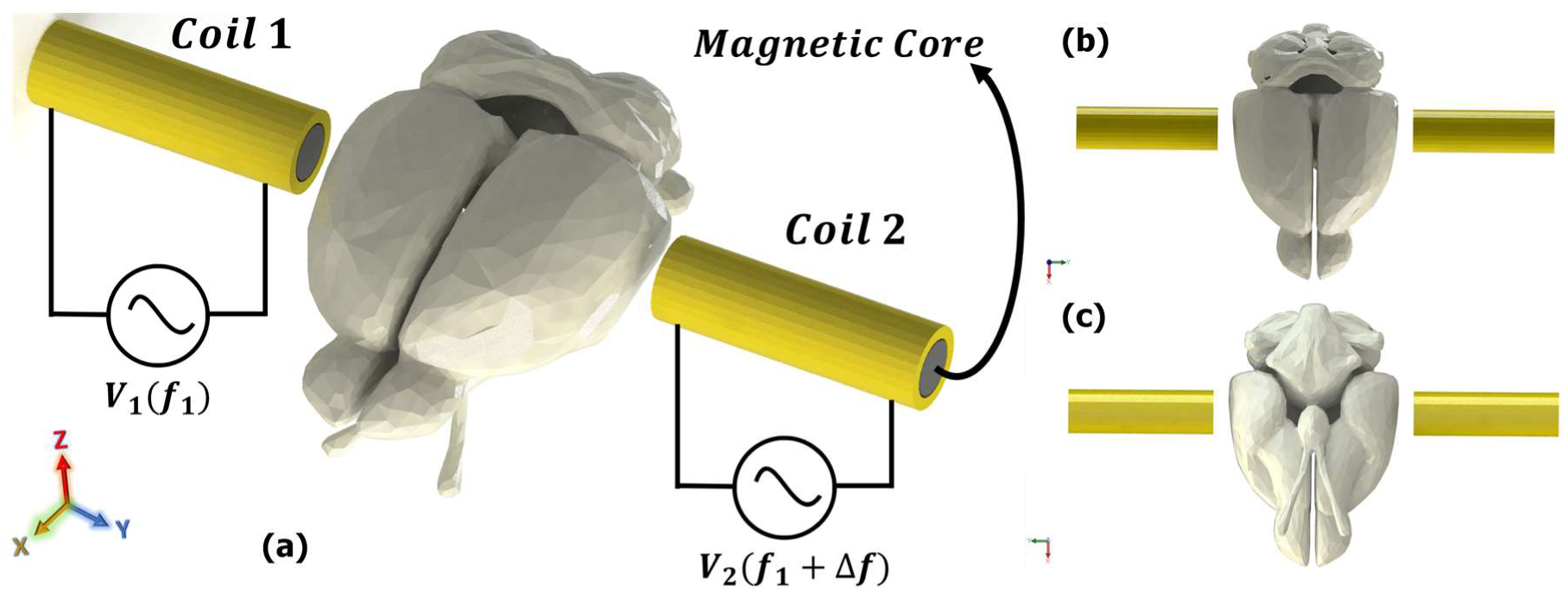
(a) The 3D rat brain model and MTI coils’ configuration used in COMSOL simulations. Coils 1 and 2 are excited using two different sources with the same amplitude but slightly different frequency Δ*f*. The coils have MnZn magnetic core with relative permeability of 17 that boosts the generated magnetic field. (b) and (c) show the top and bottom view of rat brain model, respectively.

**Fig. 3.**
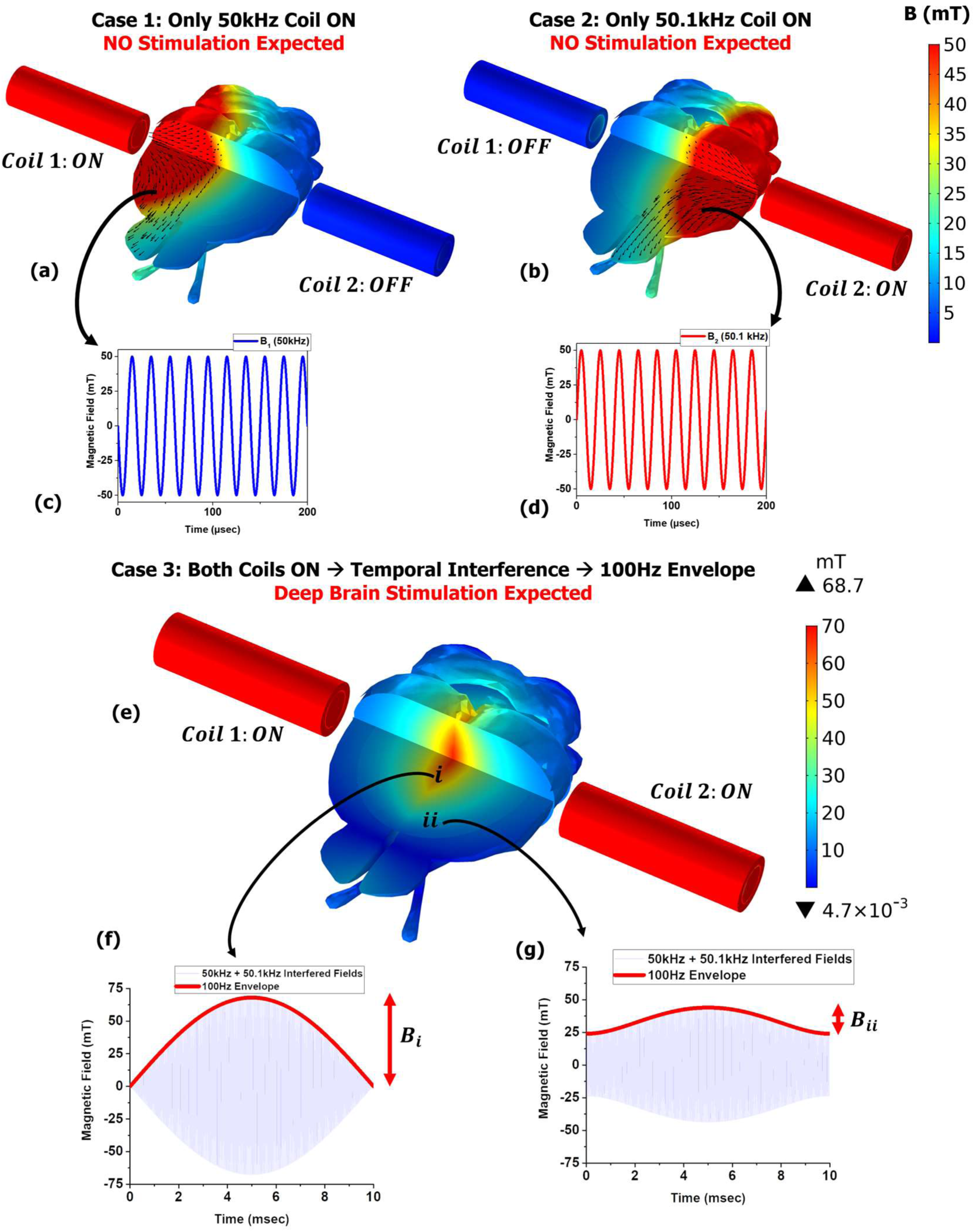
(a) Magnetic field distribution inside the cross-sectioned rat brain when coil 1 is ON and coil 2 is OFF. Here the brain is only impacted by 50kHz high frequency field which cannot stimulate the brain; (b) magnetic field distribution inside the rat brain when coil 1 is OFF and coil 2 is ON. Here the brain is only impacted by 50.1kHz high frequency field and therefore no stimulation happens; (c) and (d) the time domain 50kHz and 50.1kHz Sine wave magnetic field produced by coil 1 and coil 2, respectively; (e) The amplitude of 100Hz low frequency envelope formed as a result of temporal interference of fields from coil 1 and 2 when both coils are ON; (f) and (g) show the time domain magnetic field signal in central and peripheral region when both coils are ON. The amplitude of low frequency envelope is large at the center and small in peripheral regions.

### Field strength validation of MTI technique for neural stimulation

In order to activate neurons, the gradient of induced electric field needs to be larger than a certain threshold. The threshold of the gradient of electric field is about 11k V/m^2^ [12, 13]. Fig. 4a-4d show the simulation results for the generated electric field and E-field gradient by MTI coils. As we can see the induced electric field (E_x_) and electric field gradient (dE_x_/dy) are focused in central part of the brain with a high spatial resolution. The electric field gradient is maximum in the center and is as high as 16k V/m^2^, which is higher than the threshold value 11kV/m^2^. The stimulated region in deep brain area, where the E-field gradient is above the threshold value, is around 2∼3mm, which indicates that MTI has a very high spatial resolution. Fig. 4e shows the induced electric field gradient of repetitive MTI (rMTI) where we apply a gradient pulse every second to the deep brain region. Repetitive transcranial magnetic stimulation has been extensively researched in the past two decades and it has been widely shown that this technique can be very effective for the treatment of major depression disorder, Parkinson’s disease, and stroke [14, 15, 16, 17]. Fig. 4f and 4g show the induced electric field and E-field gradient along the y-axis line [shown in Fig. 4a] at different career frequencies *f*_1_. As it is shown the induced fields linearly increase with increasing the frequency, a phenomenon that is due to the Faraday’s law of electromagnetic where the induced electric field is linearly proportional to the rate of change of the magnetic field over time. This phenomenon can be a great upper hand for MTI technique where we can achieve a larger induced electric field and field gradient just by increasing the career frequencies *f*_1_ and *f*_2_, without the need to increase the current which can significantly increase the heating in the system. Spatial resolution of 2∼3mm that was discussed before can be further improved by reducing the career frequencies or the current in a fashion that only the desired area at the center is impacted by the field gradients larger than threshold value. In Fig. 4g, for instance, at career frequency 40kHz, a much smaller area—around 1mm— is impacted by field gradients greater than threshold value. In other word, by carefully setting the career frequency we can further improve the spatial resolution and only stimulate a very small area deep inside the brain. Fig. 4h shows the time domain electric field at the center of the brain at different career frequencies *f*_1_. The low frequency envelope shaped by the temporal interference of two career frequencies are shown in different colors. As it was discussed before, the induced electric field linearly increases by increasing the career frequencies, while the low frequency envelope of 100Hz is still preserved at all cases.

**Fig. 4.**
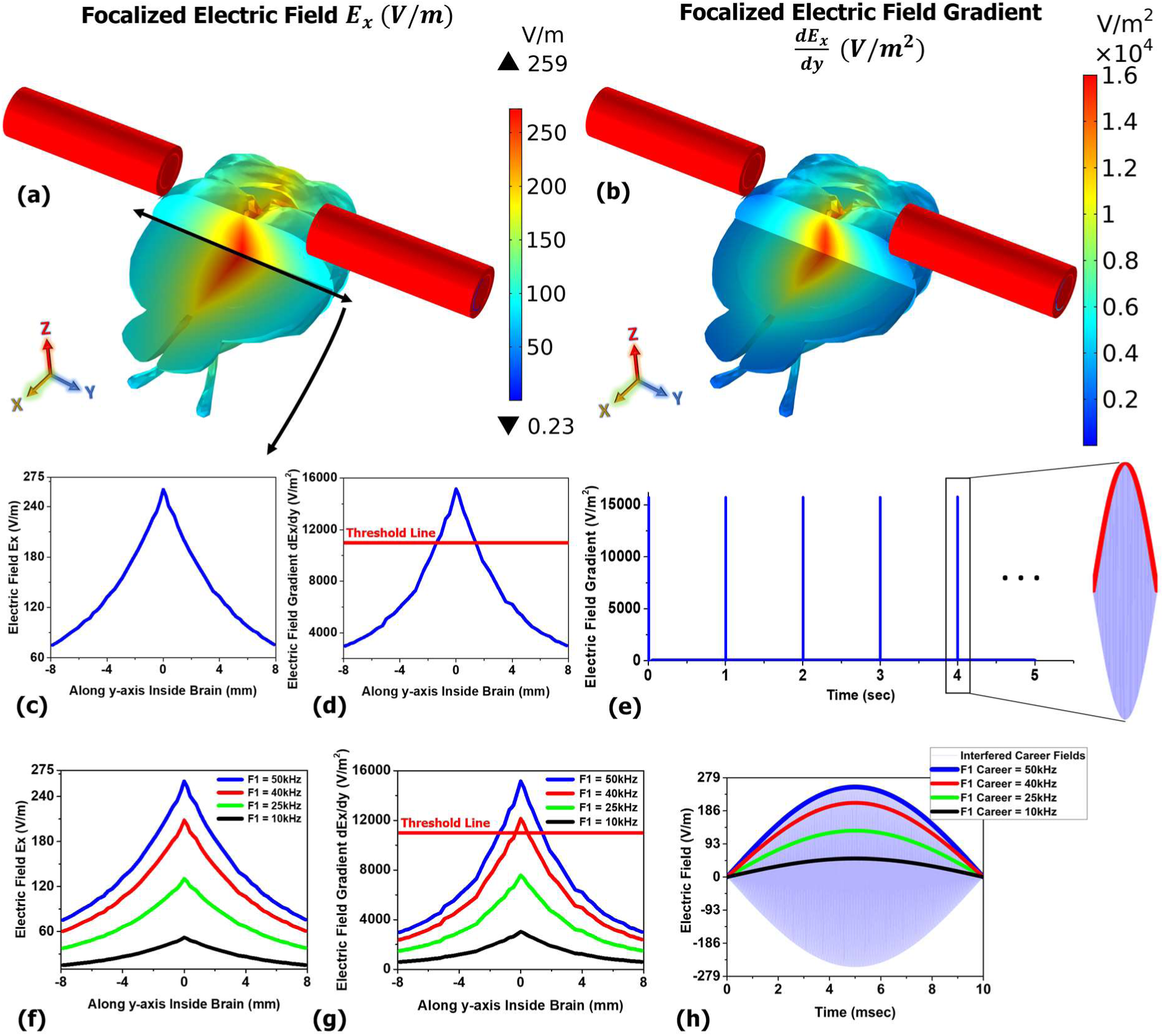
(a) and (b) show the induced electric field and E-field gradient, respectively, both focused in deep brain area with a high resolution; (c) and (d) show the E-field and E-field gradient along the y-axis line shown in fig. 4a. As we can see the induced E-field gradient at the center is above the threshold value 11kV/m^2^; (e) The induced E-field gradient of repetitive MTI (rMTI) where a gradient pulse is applied to the brain every second; (f) and (g) show the induced electric field and E-field gradient along the y-axis line [shown in Fig. 4a] at different career frequencies *f*_1_. As it is shown the induced fields linearly increase with increasing the frequency, a phenomenon that is due to the Faraday’s law of electromagnetic; (h) shows the time domain electric field at the center of the brain at different career frequencies *f*_1_. As it was discussed before, the induced electric field linearly increases by increasing the career frequencies, while the low frequency envelope of 100Hz is still preserved at all cases.

### Moving the focal point of MTI by changing the applied currents to the coils

In previous parts we investigated the MTI technique for focusing a magnetic or electric beam deep at the center of the brain. It was shown that using two MTI coils we can successfully focalize the low frequency field component precisely at the center. In this section we will discuss about how to focalize the field at different depths inside the brain, and not necessarily at the center. MTI technique allows us to focalize the beam at any depth inside the brain just by adjusting the ratio of the currents applied to the coil 1 and 2. The equation (eq. 1) for calculating the amplitude of low frequency component is shown in the next experimental section. According to this formula, maximum low frequency amplitude occurs where the amplitude of high frequency electric (or magnetic) fields *E*_1_(*f*_1_) and *E*_2_(*f*_1_ + Δ*f*) are equal. When applied current to both coils are equal, shown in Fig. 5a, the field distribution *E*_1_(*f*_1_) and *E*_2_(*f*_1_ + Δ*f*) generated by the coils on two sides of the brain are symmetric and therefore they have equal value at the center of the brain. That is why the focal point is maximum at the central region. When one of the coils is excited at a larger current, however, the focalized beam is no longer at the center and will be shifted towards the coil with weaker current. In Fig. 5b, for instance, where applied current to coil 2 is two times as large as applied current to coil 1, the beam is shifted towards coil 1; this is because coil 1 produces a weaker electric field compared to coil 2, and therefore the location where both high frequency electric fields are equal—in other word, the location where beam focalization happens—is in the region close to coil 1. Fig. 5c shows the simulation results where applied current to coil 2 is four times as large as applied current to coil 1. The focalized beam in this case is further shifted towards coil 1 compared to Fig. 5b and 5a. Fig. 5d shows the normalized electric field along y-axis inside the brain at different current ratios applied to coil 1 and coil 2. As it was discussed, by changing the ratio of the current 1 and 2 we can readily adjust the focal depth of MTI technique and shift the focal point towards left or right, without changing the location of the coils or readjusting the setup.

**Fig. 5.**
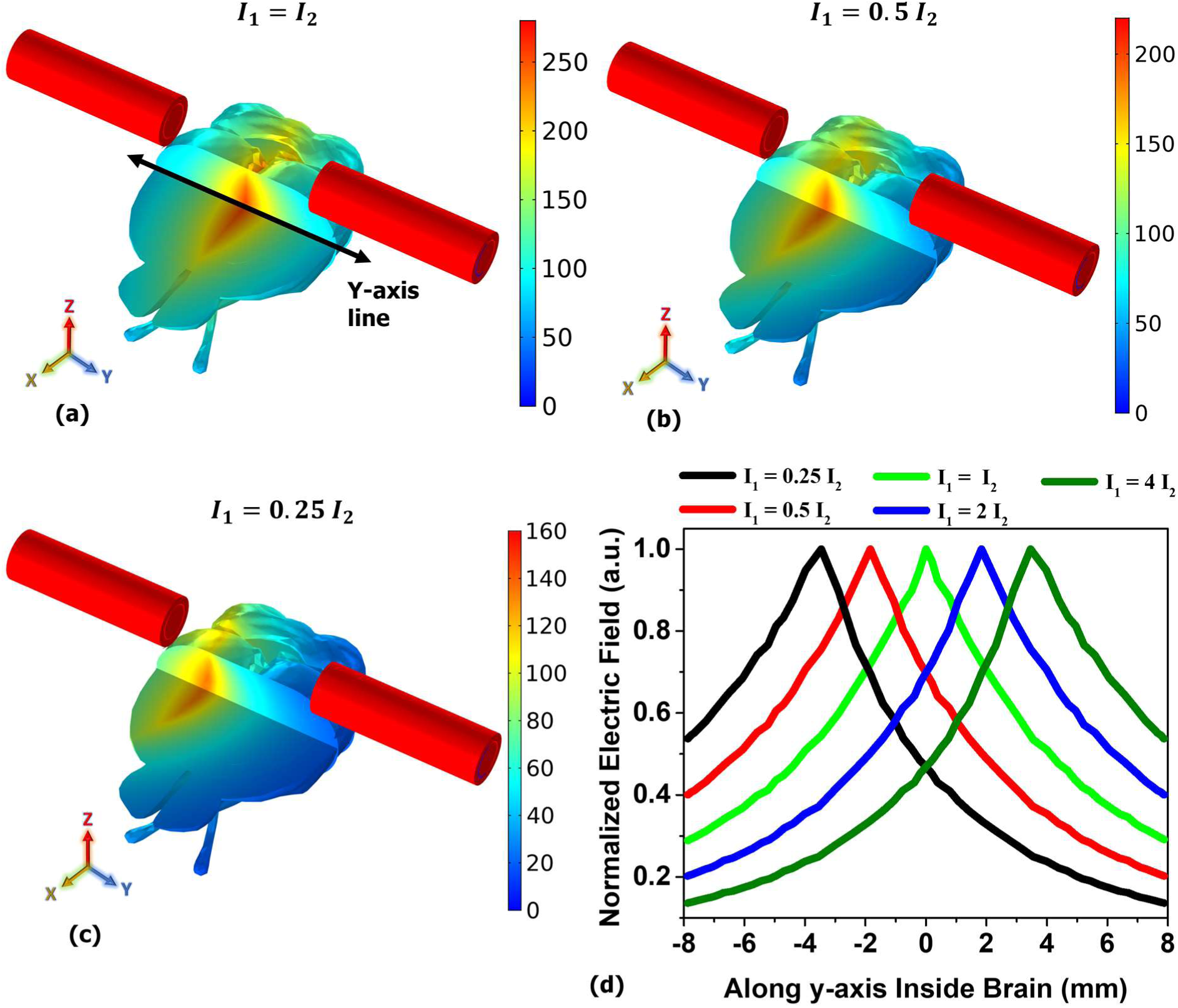
(a) electric field distribution when applied current to coil 1 and coil 2 are equal. As it is shown the beam focalization is symmetric and MTI beam is maximum at the center; (b) and (c) beam focalization where applied current to coil 2 is two times and four times as large as applied current to coil 1, respectively. As it is shown, by changing the ratio of applied current to the MTI coils the focalized beam can be shifted towards the coil with weaker current. In these two figures the focal point is shifted towards coil 1 (left side); (d) normalized electric field along y-axis inside the brain at different current ratios applied to coil 1 and coil 2. As it is shown we can readily shift the focal point towards left or right just by changing the ratio of the current, without changing the location of the coils or readjusting the setup.

### Experimental demonstration of magnetic temporal interference concept

In previous sections the theory, capabilities, and advantages of MTI technique were discussed based on COMSOL simulation results. In this section, we will experimentally investigate the concept of MTI technique and show that two fields with slightly different frequencies and generated by two different sources can focus a magnetic field beam at the central region between the coils. In this experiment we use two home-made solenoid coils with 2cm dimeter, 2cm height, and 15 turns. A matching capacitor is also added in series to each coil to cancel the inductance value and to assure that series LC circuit is in resonance condition at 50kHz operational frequency. Putting the circuit in resonant mode significantly reduces the reflection power and helps to achieve maximum magnetic field from the coils. We excite coil 1 at 50kHz and coil 2 at 50.1kHz, both under a fixed current of 1Amp. The we place the two coils on the two sides of a circular region with radius of 10cm, which corresponds to the radius of human brain model used in Fig. 1. This circular region is shown in Fig. 6a, where we have an array of 21×21 measurement data points, or 441 pixels in total. After exciting the coils under mentioned conditions, we measure the amplitude of the 50kHz and 50.1kHz magnetic fields at each pixel using a 1mm size home-made search coil connected to the spectrum analyzer. Therefore, at each pixel, we have two data points: FFT amplitude of magnetic field at *B*_1_(*f*_1_) and *B*_2_(*f*_1_ + Δ*f*). Then using the standard formula below we can calculate the amplitude of the low frequency envelope:

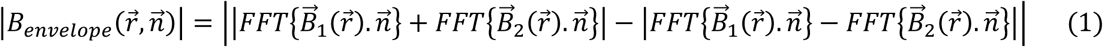

**Fig. 6.**
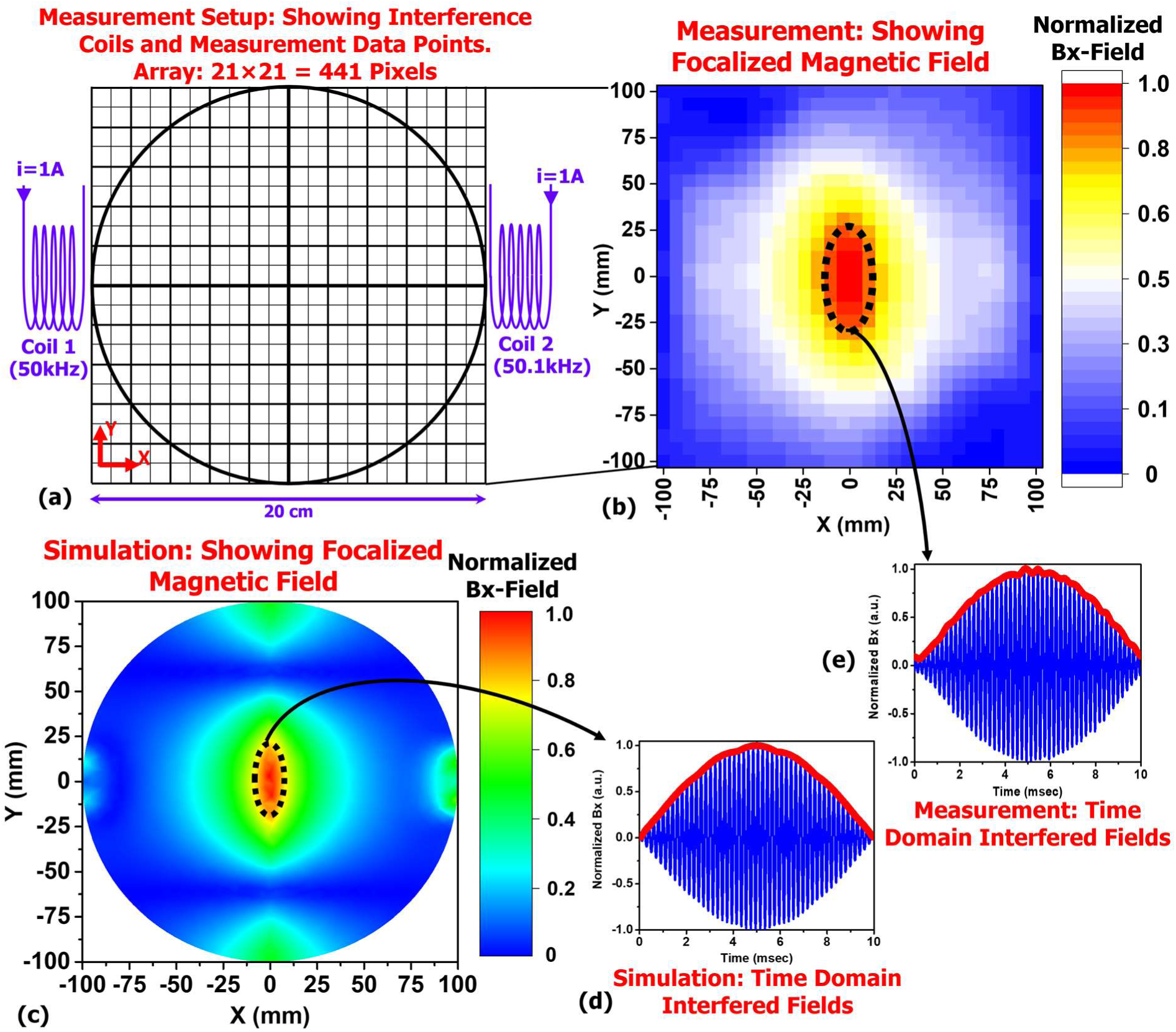
(a) The diagram of experimental setup used to verify the temporal interference concept. The region between two coils include an array of 21×21 measurement data points, or 441 pixels in total. coil 1 and 2 are excited at 50kHz and 50.1kHz, respectively, both under a fixed current of 1Amp; (b) the 2D measurement results of the normalized low frequency magnetic field envelope in the region between two coils. it is shown the low frequency magnetic beam is focused at the central region between two coils.; (c) COMSOL simulation results of the same set up, coils, and the region size. The simulation results confirm the measurement data and show that the magnetic beam is maximum in the central region; (d) and (e) show the time domain measurement and simulation results, respectively, at the central region. The plots include the interfered fields and low frequency envelope shown in red.

Based on the simulation results from previous sections we already know that the magnetic field low frequency envelope is maximum along x-axis, which is the axis parallel to the solenoid axis, and other field components along y- and z-axis are significantly smaller in comparison. Thus, by carefully aligning our search coil along x-axis we only pick up the fields *B*_1_(*f*_1_) and *B*_2_(*f*_1_ + Δ*f*) along this axis and then calculate the low frequency envelope Bx using equation (1). It is notable that even though this measurement is done in the air medium, it is expected the temporal interference of the two fields will lead to the same results and same focality even in a biological medium. This is because (1) the measurement region and its size are several orders of magnitude smaller than the operational wavelength (∼6000m), and this means that field distribution is completely inductive. If, for instance, the operational frequency were extremely high and therefore the wavelength was comparable to our measurement area, then the field distribution might be on propagation mode and therefore temporal interference pattern and focality can be significantly impacted by adding a biological tissue or a high conductivity material close to the coils; (2) operational frequency is extremely low and this means that electrical conductivity of the biological tissue, e.g. brain tissue, is very close to that of air. A small electrical conductivity reduces the power dissipation in the tissue, as well as reduces the magnetic field distortion in the medium. Therefore, we can safely say that the magnetic field temporal interference results at this frequency range in the air should be a good representation of the results in a biological medium such as human brain. Fig. 6b shows the 2D measurement results of the normalized amplitude of the low frequency magnetic field envelope in the region between two coils. As it is shown the low frequency magnetic beam is focused at the central region between two coils. Fig. 6c shows the COMSOL simulation results of the same set up, coils, and the region size. The simulation results confirm the measurement data and show that the amplitude of low frequency magnetic field envelope is maximum at the center. Fig. 6d and 6e show the time domain measurement and simulation results, respectively, at the central region which include the interfered fields and low frequency envelope shown in red. Outside the central region the interference is partial and low frequency envelope is smaller than the value shown in these figures.

## Conclusion

We have proposed a novel brain stimulation technique that works based on temporal interference of two high-frequency magnetic fields. This technique could be applicable to non-invasive, high-resolution, and focalized deep brain stimulation. We showed that a magnetic or electric field beam can be non-invasively focused at any depth inside the brain. The induced focalized electric field inside the brain can be enhanced by increasing career frequencies applied to the two electromagnetic coils while keeping the temporally interfered envelope at a fixed value such as 100Hz. In addition, the focal point inside the brain can be readily moved by changing the ratio of the currents applied to the two coils, or by readjusting the position of the coils with respect to the brain. We also experimentally demonstrated the concept of temporal magnetic interference and showed that the low-frequency magnetic field envelope can be focused at a point between the two coils.

